# Neural Sensitivity to Word Frequency Modulated by Morphological Structure: Univariate and Multivariate fMRI Evidence from Morphologically Complex Words

**DOI:** 10.1101/2025.11.20.689262

**Authors:** Joonwoo Kim, Solbin Lee, Kichun Nam

## Abstract

A central question in psycholinguistics in visual word recognition is whether morphologically complex words are obligatorily decomposed into stems and affixes during visual word recognition or whether whole-word access can occur when forms are frequent and familiar. The present study investigated how morphological complexity and lexical frequency jointly shape neural responses by leveraging Korean nominal inflection, whose transparent stem–suffix structure permits a clean dissociation between base (stem) frequency and surface (whole-word) frequency. Twenty-five native Korean speakers completed a rapid event-related fMRI lexical decision task involving simple and inflected nouns that varied parametrically in both frequency measures. Representational similarity analysis (RSA) revealed robust encoding of surface frequency—but not base frequency—in the inferior frontal gyrus (IFG) pars opercularis and supramarginal gyrus (SMG), with significantly stronger correlations for inflected than simple nouns. Univariate analyses converged with this result: surface frequency selectively increased activation for inflected nouns in inferior parietal regions, whereas base frequency showed no reliable effects in any ROI. These findings challenge models positing obligatory pre-lexical decomposition, instead supporting accounts in which morphological processing is shaped by post-lexical, usage-driven lexical statistics. Taken together, our findings shed light on a distributed perspective on morphological processing, suggesting that structural and statistical factors jointly constrain access to morphologically complex forms.

## INTRODUCTION

Human language comprehension requires efficient recognition of morphologically complex words. A central question in psycholinguistics in visual word recognition is whether the cognitive system obligatorily decomposes these words into morphemes before meaning access or whether whole-word representations can be accessed directly for familiar forms.

Evidence for obligatory decomposition comes primarily from masked-priming studies, where briefly presented primes like *hunter* facilitate recognition of *hunt*, even when semantically opaque (e.g., *corner–corn*) relative to orthographic controls (Longtin et al., 2003; Rastle et al., 2004). Such priming, with prime durations as brief as ∼50 ms, suggests rapid morpho-orthographic segmentation that precedes semantics. ERP and MEG work corroborates this view, showing early negativities before semantic divergence (Lavric et al., 2011; Lewis et al., 2011). It is important to note that the empirical and theoretical frameworks reviewed here pertain specifically to visual word recognition, where orthographic structure provides the primary cues to morphological segmentation. Phonological processing can exhibit different morphological dynamics, particularly in languages with rich morphophonology, and therefore lies outside the scope of the present investigation.

In contrast, distributional perspectives argue that processing depends on lexical statistics, transparency, and predictability (Baayen et al., 2011; Bybee, 1995; Milin et al., 2017). High-frequency or predictable forms may be accessed holistically, whereas rare or novel forms are decomposed. In lexical decision and naming tasks, surface frequency, the token frequency of the whole form, often better predicts recognition than base frequency, the cumulative frequency of words sharing a stem (Bertram et al., 2000; Kuperman et al., 2009). Morphological priming effects are likewise modulated by transparency and familiarity (Feldman et al., 2015; Jared et al., 2017). Connectionist models further show how distributional learning can yield decomposition-like effects without an explicit parsing mechanism (Plaut & Gonnerman, 2000). Together, these findings suggest morphological processing may reflect both automatic segmentation and flexible, frequency-driven representations.

Another critical distinction in morphological processing concerns the timing of decomposition relative to lexical access. Pre-lexical decomposition models propose that complex words are obligatorily parsed into constituent morphemes before whole-word meaning is accessed (Rastle et al., 2004; Taft & Forster, 1975). Under this view, recognition of *hunter* necessarily involves early access to *hunt* + *-er*, regardless of whole-word frequency. In contrast, post-lexical decomposition accounts propose that morphological structure is analyzed *after* initial whole-word recognition (Giraudo & Grainger, 2000, 2001). Here, frequent derived forms like *hunter* may be accessed directly as whole words, with morphological decomposition occurring later for semantic integration or production. Critically, these accounts make different predictions about which frequency measure should modulate neural responses: pre-lexical models predict that base (stem) frequency should drive activation patterns because stems are accessed first, whereas post-lexical models predict that surface (whole-word) frequency should dominate because initial access relies on whole-word representations. Notably, surface frequency itself can be interpreted differently across frameworks: in pre-lexical models, it reflects the frequency of recombining morphemes, whereas in post-lexical models, it indexes the frequency of stored whole-word forms. The present study leverages fMRI to test these competing predictions by examining whether sustained neural activation patterns are better explained by surface or base frequency.

Neuroimaging evidence has shown mixed results regarding whether morphological processing reflects categorical decomposition or graded statistical effects. fMRI studies report that complex forms engage a left-lateralized network including the inferior frontal gyrus (IFG), inferior parietal lobule (IPL), and ventral occipitotemporal cortex. Some findings have been interpreted as supporting categorical decomposition, which showed selective IFG activations to inflected or transparent derivations in English and Finnish (Bozic et al., 2007; Lehtonen et al., 2006), and distinct activation patterns for regular versus irregular inflections in the left frontotemporal network in German and English (Beretta et al., 2003; Tyler et al., 2004). However, other work suggests that these effects reflect graded sensitivity to phonological complexity, semantic factors, and lexical statistics rather than a unique decomposition stage (Devlin et al., 2004; Joanisse & Seidenberg, 2005). When phonological complexity is controlled, categorical differences between regular and irregular forms often disappear (Desai et al., 2006), and when differences do emerge, they are modulated by semantic variables such as semantic transparency (Bozic et al., 2013), suggesting continuous rather than discrete processing mechanisms. While much of this work has focused on the regular versus irregular distinction in verb inflection, a complementary question remains: at what level—whole-word or stem—does the brain encode lexical frequency during morphological processing?

Findings on word frequency also remain inconsistent, with some studies reporting stronger activation for low-frequency words (Fiebach et al., 2002; Fiez et al., 1999), and others showing divergent patterns depending on task and subregion (Carreiras et al., 2009; Graves et al., 2010). Critically, few studies have manipulated base frequency directly, and most report limited independent effects once surface frequency is controlled (Lehtonen et al., 2006; Pliatsikas et al., 2014), leaving unresolved whether the brain encodes stem-level or whole-word statistics during morphological processing.

Korean morphology encompasses derivation, compounding, and inflection, but these subtypes differ substantially in transparency and semantic complexity. Prior work has examined derivational or compound morphology, which can involve semantic shifts or idiosyncratic affix behavior (Kim, Lee, et al., 2025). The present study focuses specifically on nominal inflection, where invariant stems combine transparently with case and topic markers. Korean inflectional nouns have transparent stem–suffix structure and invariant stems, which together provide a useful testing ground for understanding how morphological structure and lexical statistics contribute to visual word recognition. Previous fMRI work shows that Korean readers are sensitive to structural aspect of inflection, with effects predominantly centered at the left IFG, consistent with the findings in Indo-European languages. For instance, inflected verbs that impose greater morpho-syntactic demands elicited stronger inferior frontal activation relative to inflected nouns, and violations of morphological category recruited left IFG subregions (Kim, Pyun, et al., 2025; Kim & Nam, 2025). Studies directly comparing regular and irregular verb inflections also reported equivalent level of activation in left IFG (Kim et al., 2024; Yim et al., 2006).

On the other hand, emerging evidence suggests continuous sensitivity to lexical–semantic and distributional factors. Kim et al. (2024) observed graded increases in occipitotemporal activity for regular inflections relative to uninflected and irregular forms, with lexical-semantic ambiguity (i.e., homonymy) further modulating activation in both occipitotemporal and angular gyrus regions. In the temporal domain, ERP evidence from Korean inflected verbs showed that both form-based and lexical-semantic properties emerge rapidly during visual word recognition (Kim et al., 2022). They showed that whole-word frequency modulates early ERP components (N100/N250) and interacts with stem length at N400. High-frequency inflected verbs were accessed more holistically, whereas lower-frequency items showed stronger stem-length effects, a pattern consistent with frequency-contingent decomposition rather than uniform pre-lexical parsing.

Together, these findings suggest that, in Korean as in Indo-European languages, readers are sensitive to morphological structure, yet sustained neural responses are primarily shaped by post-lexical, usage-driven statistics. This pattern raises an important question about the representational level at which frequency information is encoded during the processing of inflectional morphology: whether neural responses track stem-based statistics, as predicted by pre-lexical decomposition accounts, or whole-word surface frequency, as predicted by post-lexical access models. To the best of our knowledge, no fMRI study has directly contrasted surface and base frequency in Korean nominal inflection, despite the transparent agglutinative structure that allows these frequency measures to be clearly dissociated.

Recent multivariate approaches offer powerful tools for examining how linguistic information is represented in the brain. Representational similarity analysis (RSA) relates neural dissimilarities to models derived from lexical statistics (Haxby et al., 2001; Kriegeskorte, 2008), enabling examination of graded representational structure. RSA has revealed fine-grained representations of syntactic, semantic, and sublexical features across languages (Bruffaerts et al., 2013; Fischer-Baum et al., 2017; Yue & Martin, 2021), allowing us to directly examine whether neural patterns reflect surface frequency, base frequency, or morphological complexity.

The present study employed a rapid event-related fMRI design with a lexical decision task to test whether brain activation patterns track morphological structure or statistical frequency. Participants judged simple and inflected Korean nouns varying in both measures. Korean provides a particularly suitable test case for several reasons. Its agglutinative morphology yields transparent stem–suffix structure, stems remain invariant across inflections, and orthographic conventions (i.e., spacing and syllable blocking) clearly indicate morphemic boundaries, all of which permit precise frequency estimation. Crucially, inflected forms exhibit a natural dissociation between surface frequency (i.e., the frequency of the whole form) and base frequency (i.e., the frequency of the bare stem). For example, the stem 까닭 /kkadak/ “reason” can combine with different particles (e.g., 까닭은, 까닭이, 까닭을); each inflected form has its own surface frequency, but they all share the same base frequency — the cumulative count of the stem 까닭. By contrast, simple (monomorphemic) nouns such as 옥수수 /oksusu/ “corn” have no inflectional suffix, so the word form and the stem are identical and the two frequency measures necessarily converge.

We applied complementary approaches—RSA and univariate mixed-effects models—to determine whether neural responses to inflected Korean nouns primarily reflect pre-lexical decomposition or post-lexical morphemic activation. If morphological parsing occurs at the pre-lexical stage, then neural activation patterns should be more strongly explained by base frequency, which indexes the frequency of accessing the stem independent of surface form. In contrast, if inflected nouns are accessed primarily as whole-word units, consistent with post-lexical accounts, then surface frequency should provide a better account of both amplitude-based responses and the representational geometry captured by RSA. The present study tests whether neural patterns align more closely with stem-driven decomposition or whole-word-driven statistical access by directly contrasting surface and base frequency in Korean nominal inflection.

## METHODS

### Participants

Twenty-five healthy adults (15 female; age: 26 ± 3.83 years) recruited from Korea University participated in the experiment. They were all native Korean speakers, right-handed (confirmed by the scores on the Edinburgh Handedness Test (Oldfield, 1971): *M* = 7.76, *SD* = 0.28) and had normal or corrected-to-normal vision. Participants were informed and provided written consent prior to the experiment and were monetarily compensated for their participation. One participant withdrew before completing the experiment, and two others were excluded due to poor behavioral performance (accuracy < 60%). One additional participant completed the experiment with adequate behavioral performance, but their fMRI data could not be analyzed due to file corruption resulting from hardware failure. Behavioral data from this participant were retained. Final sample sizes were N = 22 (11 female; age: 26.3 ± 3.97 years) for behavioral analyses and N = 21 (11 female; age: 26.4 ± 4.07 years) for all neuroimaging analyses. The sample size was determined based on prior fMRI studies using similar multivariate and univariate ROI-based approaches in the language domain (Bozic et al., 2007; Carota et al., 2016; Kim et al., 2024), which typically report medium-to-large effects in comparable tasks with sample sizes ranging from 16 to 24 participants. Given the within-subjects design and the use of trial-wise beta estimation, this sample size affords adequate power to detect region-level effects of morphology and lexical frequency using both traditional and multivariate analyses. The study was conducted in accordance with the Declaration of Helsinki and approved by the Institutional Review Board of Korea University (protocol code KUIRB-2021-0427-01, 21 December 2021).

### Stimuli

A total of 120 Korean nouns were selected from the Korean Sejong corpus (Kang & Kim, 2009), which contains approximately 15 million words. Korean is an agglutinative language with a transparent morphological system, in which grammatical relations are marked by attaching case particles or topic markers to noun stems. Furthermore, the Korean writing system (Hangul) encodes syllables both orthographically and phonologically, enabling precise control of sublexical units. These features make Korean particularly well-suited for isolating effects of morphological complexity, stem-level access, and syllable-level frequency during visual word recognition.

Morphological complexity was orthogonally manipulated such that half of the stimuli were morphologically simple nouns, and the other half were inflected nouns. Each condition included an equal number of bi-syllabic and tri-syllabic words, ensuring that word length was fully balanced across conditions. Simple nouns consisted of bi- or tri-syllabic stems, while inflected nouns were composed of mono- or bi-syllabic stems followed by a monosyllabic inflectional suffix (e.g., case or topic marker). For example, 숭어 (trout) and 뱀은 (snake + TOPIC) illustrate the contrast between simple and inflected forms, with 옥수수 (corn) and 까닭은 (reason + TOPIC) showing the contrast between a tri-syllabic simple noun and a tri-syllabic inflected noun consisting of a bi-syllabic stem (까닭) with the topic particle -은. The inflected nouns were formed using three common Korean case/topic particles. Their distribution was as follows: accusative –을/–를 (n = 29, 48.3%), topic –은/–는 (n = 21, 35.0%), and nominative –이/–가 (n = 10, 16.7%). This distribution closely aligns with the natural frequency distribution of case and topic in the given corpus (Kang & Kim, 2009).

Word frequency was operationalized using Zipf-measure (van Heuven et al., 2014), a log-transformed word frequency per billion words in the given corpus. Stem frequency was similarly transformed using the same Zipf-measure to allow for direct comparison across lexical levels. **Fig. 1** illustrates the frequency distributions for all words (**Fig. 1A**), simple nouns (**Fig. 1B**), and inflected nouns (**Fig. 1C**), with descriptive statistics summarized in **Table 1**.

**Figure 1.**
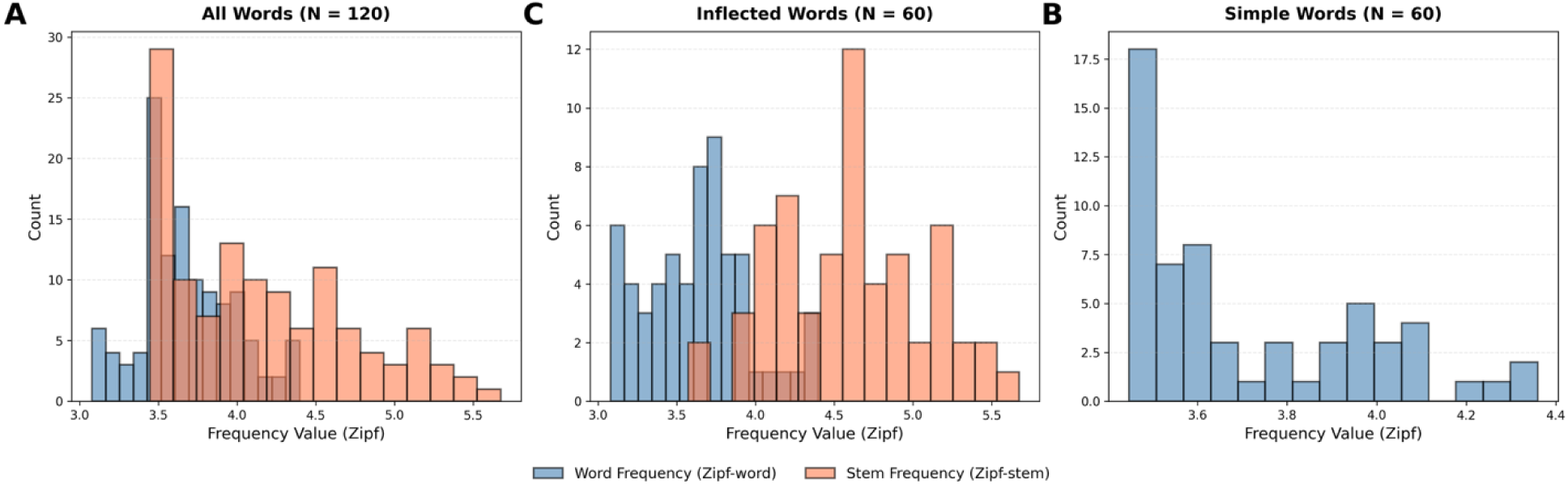
Frequency distributions across morphological conditions. (A) All words (N = 120) show overlapping distributions for surface frequency (blue) and base frequency (red). (B) Inflected nouns (N = 60) show distinct distributions for surface and base frequency measures. Histograms use 15 bins; shaded regions indicate overlapping frequencies. (C) Simple nouns (N = 60) display only surface frequency, as surface equals base by definition for monomorphemic words.

**Table 1.** Descriptive statistics of the stimuli set.

| Condition | Word length | # Morphemes | Surface freq | Base freq | 1st syllable freq |
| --- | --- | --- | --- | --- | --- |
| Simple<br>(*N* = 60) | 2.5<br>(0) | 1<br>(0) | 3.72<br>(0.27) | 3.72<br>(0.27) | 2.34<br>(0.45) |
| Inflected<br>(*N* = 60) | 2.5<br>(0) | 2<br>(0) | 3.63<br>(0.33) | 4.58<br>(0.48) | 2.25<br>(0.48) |
| Statistical<br>comparisons | - | - | *t* (59) = 1.62<br>*p* = .111 | *t* (59) = -12.32***<br>*p* < .001 | *t* (59) = 1.16<br>*p* = .251 |

*Note.* Word length = Word length in syllable; # Morphemes = Number of morphemes; Surface freq = Zipf-scale word frequency; Base freq = Zipf-scale stem frequency; 1st syllable freq = log-transformed first syllable frequency per million. Values in parentheses represent standard deviations.

Importantly, word frequency (Zipf) was matched across simple and inflected conditions (*t*(59) = 1.62; *p* = .111), whereas stem frequency (Zipf-stem) was not explicitly controlled. This was due to the structural properties of Korean: in simple nouns, the stem and surface word are identical, yielding the same frequency value. In contrast, inflected nouns contain an additional morpheme, such that the word frequency reflects the full surface form, while the stem frequency corresponds to the base noun. Given this natural divergence, Zipf and Zipf-stem were modeled separately in all statistical analyses, allowing us to dissociate effects of surface-level lexical access and stem-level frequency. To validate the dissociation between these two measures, we examined their distributional properties. Base frequency exhibited greater dispersion than surface frequency across all stimuli (coefficient of variation [CV]: 0.141 vs. 0.081) and within inflected nouns (CV: 0.103 vs. 0.090), a difference confirmed by Levene’s tests (all words: *F* = 47.22, *p* < .001; inflected only: *F* = 7.23, *p* = .008). For inflected nouns, the two measures were moderately correlated at the item level (Pearson *r* = .60, *p* < .001), yet their representational similarity structures were only weakly related (Spearman ρ = .24, *p* < .001), indicating that surface and stem frequency cannot be treated as interchangeable. Additional representational-similarity validation is provided in **Fig. S1**. Although stem syllable counts necessarily differ between conditions (simple nouns: 2–3 syllable stems; inflected nouns: 1–2 syllable stems + monosyllabic suffix), total word length was fully balanced across conditions, reducing the likelihood that sublexical length differences at the stem level confound the frequency effects reported here. Finally, first-syllable frequency was also matched across conditions (*t*(59) = 1.16, *p* = .251).

The same number of pseudowords (*N* = 120) were created to serve as fillers in the lexical decision task. All pseudowords were manually generated by recombining syllable blocks drawn from real Korean noun stems and inflectional suffixes. Each pseudoword was constructed to be orthographically and phonologically legal in Korean while carrying no meaning, as verified against the Standard Korean Dictionary. These pseudowords matched the real word stimuli in both form and length (i.e., 2–3 syllable) and were evenly divided into morphologically simple and inflected forms. Inflected pseudowords consisted of a real suffix combined with a pseudo-stem while simple pseudowords contained only a pseudo-stem. All pseudowords were generated by randomly combining syllables from real word stems and suffixes, which were orthographically and phonologically legal but none of the pseudowords carried meaning according to the Standard Korean Dictionary.

Finally, 30 non-linguistic masks, consisting of a string of asterisks (*****), were used as a baseline condition for a comparison with experimental conditions. These masks were visually matched to word stimuli in font and size, but contained no linguistic or orthographic information. Participants responded to these items using the same lexical-decision buttons, allowing us to estimate low-level visual and motor activation while controlling for perceptual properties of lexical stimuli.

### Procedure

A rapid event-related fMRI design was employed using a visual lexical decision task. Participants were instructed to respond, as quickly and accurately as possible, whether each presented stimulus is a real word or not. Each trial began with a fixation cross (+) displayed for 100 ms, followed by a target stimulus presented at the center of the screen for 1000 ms. After pressing response button, blank screen was shown during the inter-trial interval (ITI), which was jittered with an average duration of 4000 ms (range: 1000–7000 ms). Stimuli were presented in a pseudo-random order that included words, pseudowords, and a non-linguistic baseline mask. The trial order and ITI durations were optimized using optseq2 (https://www.nitrc.org/projects/optseq). All visual stimuli were presented in white Courier New font (size 34 pt) on a black background, controlled using E-Prime 2.0 Professional (Psychology Software Tools, Pittsburgh, PA). Prior to the main experiment, participants completed a practice session consisting of 12 randomly ordered trials. Participants proceeded to the main task only if they achieved at least 70% accuracy during practice.

### Image acquisition and preprocessing

MRI data were acquired at the Korea University Brain Imaging Center using a Siemens Magnetom Trio 3T MRI scanner (Erlangen, Germany). Functional images were acquired using T2*-weighted gradient EPI (echo-planar imaging) sequences (TR = 2000 ms; TE = 20 ms; flip angle = 90◦; field of view = 240 mm; slice thickness = 3 mm; no gap for 42 slices; matrix size = 80 × 80; and voxel size = 3 × 3 × 3 mm). A high-resolution structural image was acquired for each participant using a T1-weighted structural images were acquired with a 3D MP-RAGE (magnetization-prepared rapid gradient echo) sequence (TR = 1900 ms; TE = 2.52 ms; flip angle = 90◦; field of view = 256 ms; matrix size = 256 × 256; and voxel size = 1 × 1 × 1 mm) covering the whole head.

Preprocessing of the fMRI data was performed using SPM12 (Wellcome Centre for Human Neuroimaging, London, UK; https://www.fil.ion.ucl.ac.uk/spm/software/spm12/) in conjunction with custom-built MATLAB scripts. The first three volumes of each session were discarded to allow for magnetic field stabilization. The remaining functional images were first realigned to the first volume to correct for head motion using a six-parameter rigid-body transformation. Slice timing correction was applied to adjust for differences in acquisition timing between slices, using the middle slice as reference. Each participant’s structural image was then co-registered to the mean functional image and segmented. The deformation fields obtained during segmentation were subsequently applied to normalize the functional images to the standard MNI space, with resampling to 3 × 3 × 3 mm isotropic voxels. All normalized images were smoothed with a 6 mm full-width at half-maximum (FWHM) Gaussian kernel to improve signal-to-noise ratio. The six motion parameters from the realignment step were included as nuisance covariates in all first-level general linear models (GLMs).

### Behavioral data analysis

Behavioral analyses were conducted on 22 participants who completed the task with adequate performance (accuracy ≥ 60%) and had valid behavioral data. We analyzed reaction times (RT) using linear mixed-effects models (LMMs) and accuracy (ACC) using generalized linear mixed-effects models with a binomial link. All analyses reported below were conducted on word trials only, excluding pseudowords trials. For RT, only correct trials were included, and RTs were log-transformed to reduce skewness. Separate models were constructed for whole-word frequency (Zipf-word) and stem frequency (Zipf-stem), each including Morphological Condition (Simple vs. Inflected) and their interaction as fixed effects. Random intercepts for participants and items were included to account for repeated measures (Baayen et al., 2008). Accuracy data was analyzed using a binomial logistic regression in a general linear mixed effects model. These analyses were performed in R (R Core Team, 2022) using the lme4 package (Bates et al., 2015), and the significance of fixed effects was assessed using Satterthwaite’s approximation for degrees of freedom, as implemented in the lmerTest package (Kuznetsova et al., 2017).

### Whole-brain analysis

All neuroimaging analyses were conducted on 21 participants who met behavioral criteria and had valid fMRI data (see Participants section for exclusion criteria). First-level analyses modeled four contrasts for each participant: Inflected > Simple, Inflected > Baseline, Simple > Baseline, and Word > Baseline. The Inflected and Simple contrasts included only real word trials from their respective morphological conditions. The Word > Baseline contrast collapsed across all real word trials (both Inflected and Simple conditions), excluding pseudowords. Events were modeled with durations of 2 s and convolved with the canonical hemodynamic response function (HRF), and six motion parameters were included as nuisance regressors. At the second level, one-sample *t*-tests were performed on first-level contrast images. Whole-brain results were thresholded at a voxel-wise *p* < .001 (uncorrected) with a minimum cluster size of 30 voxels, consistent with exploratory standards in prior morphological and lexical processing fMRI studies (e.g., Kim et al., 2021; Newman et al., 2010). Activation patterns were further evaluated using anatomical masks from the Harvard-Oxford Atlas.

### ROI selection

Our main analyses, including univariate contrasts and representational similarity analysis (RSA), focused on six anatomically defined, left-hemisphere regions of interest (ROIs, see **Fig. 6A**). These included two subregions of the inferior frontal gyrus (pars triangularis, pars opercularis), two temporal-occipital regions including the middle temporal gyrus (MTG), fusiform gyrus (FuG), and two regions within the inferior parietal lobule (IPL): the angular gyrus (AG) and supramarginal gyrus (SMG). ROI selection was motivated by prior literature implicating these regions in morphological and lexical-semantic processing (Binder et al., 2009; Bulut, 2022; Vigneau et al., 2006), allowing us to examine convergent and dissociable effects across multiple analytical frameworks. All ROIs were defined using the probabilistic Harvard-Oxford cortical atlas (1 mm resolution). Each region was thresholded at 25% probability and binarized and resampled to match functional image resolution (3 mm) using SPM12. All masks were visually inspected to ensure accurate anatomical localization.

### Representational Similarity Analysis (RSA)

Representational similarity analysis (RSA) was used to test whether local activation patterns reflected graded similarity based on word or stem frequency. All representational similarity analyses (RSA) were implemented using custom Python scripts in combination with scikit-learn (Pedregosa et al., 2011), SciPy (Virtanen et al., 2020), and Nilearn (Abraham et al., 2014). Analyses were performed within the six anatomically defined ROIs.

Six theoretical representational dissimilarity matrices (RDMs) were constructed based on continuous lexical frequency measures: surface-level word frequency (Zipf) and base-form stem frequency (Zipf-Stem). Each RDM encoded the absolute pairwise frequency difference between trials (i.e., Euclidean distance in a 1D lexical space), producing symmetrical matrices where larger values reflect greater dissimilarity in frequency. For each frequency measure, we created separate model RDMs for three trial subsets: (1) all word trials (**Figs. 2A, 2D**), (2) morphologically inflected nouns (**Figs. 2B, 2D**), and (3) only simple nouns (**Figs. 2C, 2F**). This yielded six theoretical RDMs in total (2 frequency models × 3 morphological conditions).

**Figure 2.**
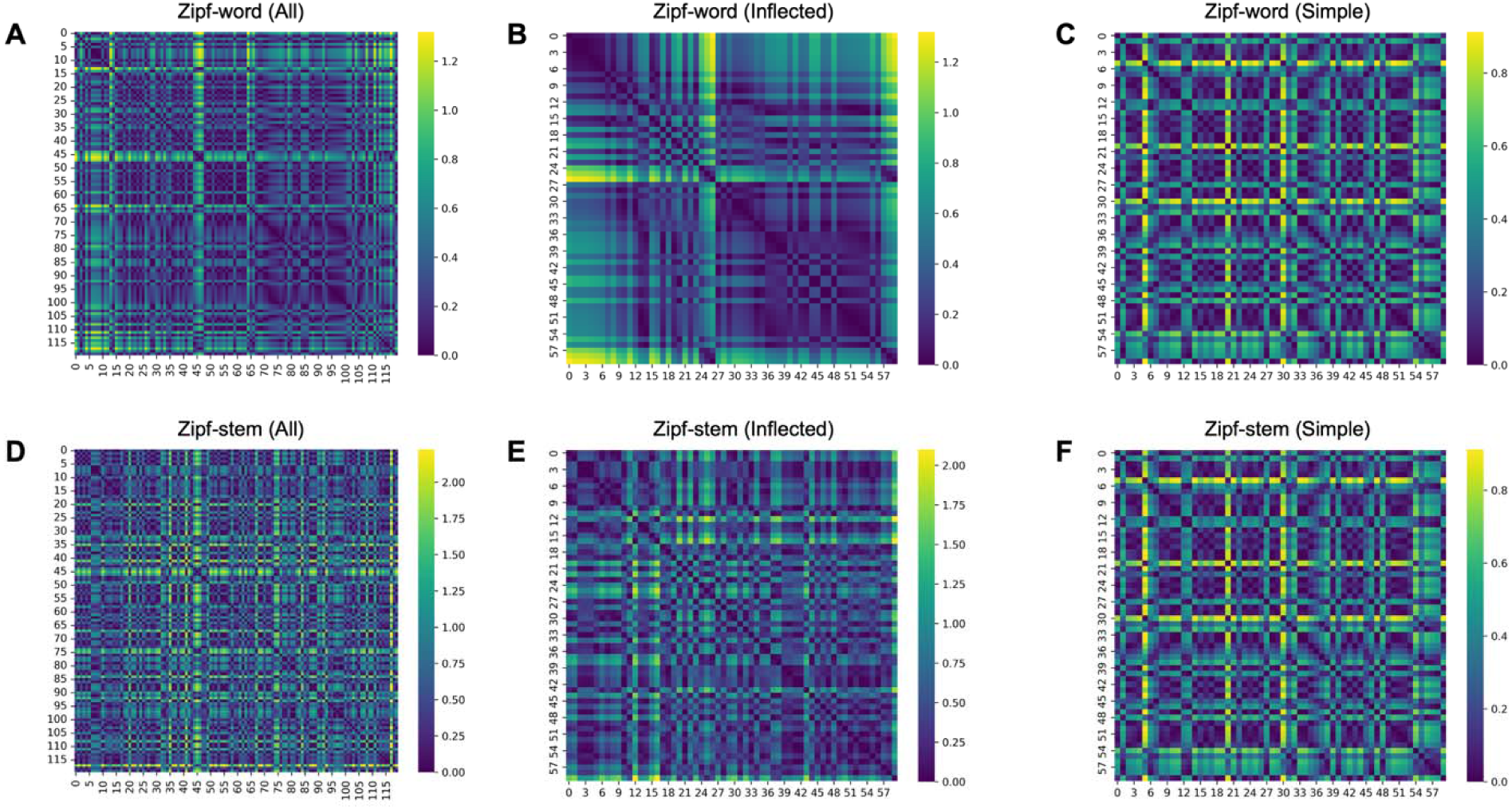
Theoretical frequency-based representational dissimilarity matrices (RDMs). Each matrix encodes the pairwise lexical distance that served as a model predictor in the RSA. Word-frequency RDMs (Zipf-word; panels A–C) and stem-frequency RDMs (Zipf-Stem; panels D–F) were computed as the absolute difference (i.e., Euclidean distance in a 1-D frequency space) in Zipf-scale frequency between two items, yielding symmetrical matrices whose values increase with greater frequency disparity. For both frequency measures we constructed three matrices: all nouns, only morphologically inflected nouns, and only simple nouns. Warmer colors denote larger frequency gaps; cooler colors denote smaller gaps.

To construct neural RDMs, we extracted trial-wise beta images per participant and ROI using NiftiMasker (standardized across voxels) and computed pairwise Pearson correlations between activation vectors for each pair of trials. Neural dissimilarity was defined as 1 minus Pearson r, resulting in a neural RDM for each subject, ROI, and condition (All, Simple, Inflected). Spearman rank correlation was used to compute the correspondence between neural and model RDMs. Correlation values were computed separately for each subject, ROI, condition, and model.

At the group level, we assessed whether neural RDMs significantly aligned with theoretical frequency structure using one-sample t-tests against zero (within-subject Fisher-z transformed Spearman values). To test whether representational structure differed across morphological conditions, we performed paired-sample *t*-tests comparing Simple vs. Inflected correlations within each ROI and frequency model. All resulting *p*-values were corrected for multiple comparisons using the false discovery rate (FDR) method (Benjamini & Hochberg, 1995).

### Univariate analysis

To examine morphological effects and lexical frequency modulations within key language-related regions, we conducted ROI-based univariate analyses. Trial-wise beta values were extracted using MarsBaR (Brett et al., 2002) and submitted to linear mixed-effects models in R.

Separate models were estimated for each frequency measure (Zipf or Zipf-stem) and ROI, with fixed effects for morphological condition (simple vs. inflected), frequency, and their interaction. Random intercepts were included for both participants and items to account for repeated measures and individual variability. These models assessed whether frequency effects varied by morphological complexity and whether either measure explained additional variance in regional activation.

## RESULTS

### Behavioral performance

**Fig. 3** presents mixed-effects model predictions for reaction time and accuracy as functions of surface (Zipf-word; **Fig. 3A**) and base frequency (Zipf-stem; **Fig. 3B**). In both the Zipf-word and Zipf-stem models for RT, morphological condition did not show a significant main effect (Zipf-word: *β* = –0.027, *SE* = 0.036, *t* = –0.85, *p* = .399; Zipf-stem: *β* = –0.028, *SE* = 0.034, *t* = –0.82, *p* = .416), nor was the interaction between morphology and frequency significant (Zipf-word: *β* = 0.009, *SE* = 0.01, *t* = –0.94, *p* = .347; Zipf-stem: *β* = 0.01, *SE* = 0.009, *t* = 1.215, *p* = .227). However, both frequency predictors showed significant main effects, with faster responses to higher-frequency words. Word frequency (Zipf-word) was associated with reduced RTs (*β* = –0.03, *SE* = 0.01, *t* = –3.08, *p* = .003; **Fig. 3A**), and the same pattern was observed for stem frequency (*β* = –0.026, *SE* = 0.009, *t* = –3.02, *p* = .003; **Fig. 3B**).

**Figure 3.**
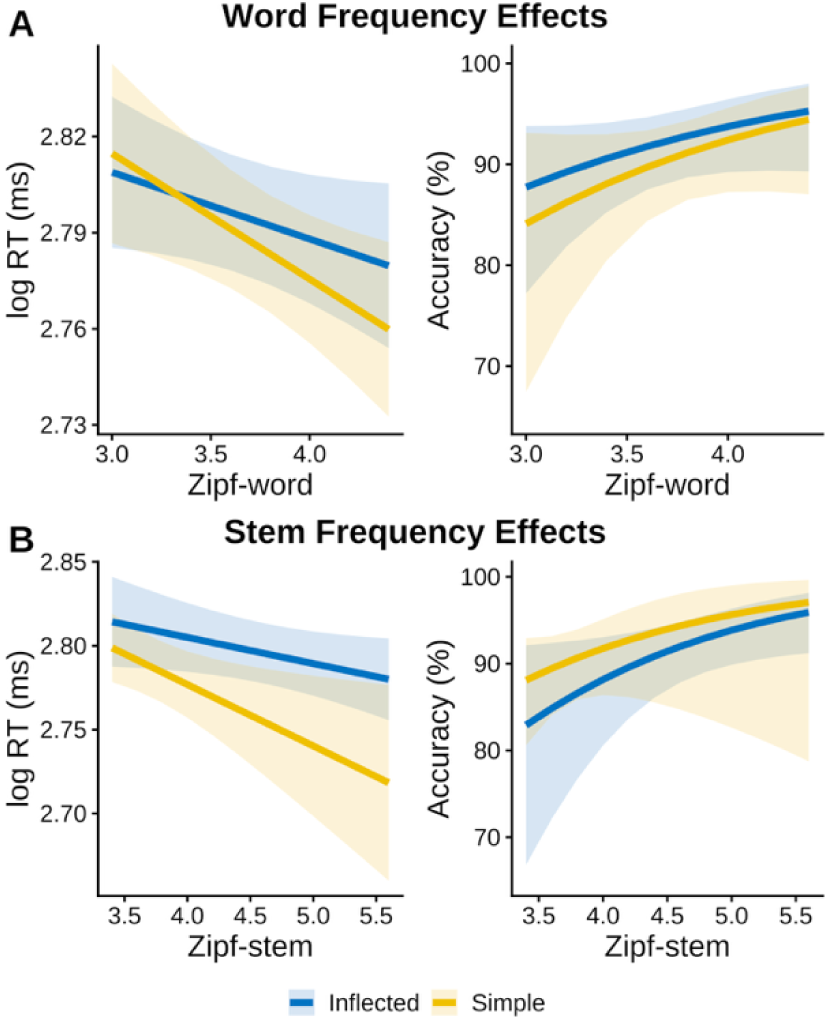
Mixed-effect model predictions for behavioral performance. Predicted reaction times (left y-axes), log-transformed milliseconds and predicted probability of correct response (right y-axes) are shown as functions of (A) word frequency (Zipf-word) and (B) stem frequency (Zipf-stem). Blue lines represent inflected nouns; yellow lines represent simple nouns; shaded ribbons indicate 95% confidence intervals (CIs).

In the accuracy model, no significant effects were found for morphological condition or its interaction with frequency (all *p*s > .83; **Fig. 3**). Both word frequency (*β* = 0.787, *SE* = 0.388, *z* = 2.03, *p* = .043; **Fig. 3A**) and stem frequency had a significant main effect (*β* = 0.699, *SE* = 0.34, *z* = 2.07, *p* = .038; **Fig. 3B**), suggesting faster latencies and greater accuracy for items with higher frequencies of either the stem or the whole-form.

### Whole-brain analysis results

**Table 2** and **Fig. 4** present the results of the univariate whole-brain analysis. All contrasts were computed at a voxel-level threshold of *p* < .001 (uncorrected) with a cluster extent threshold of *k* > 30. Anatomical labels were assigned using the Harvard-Oxford cortical atlas (HOA), and all reported clusters exceeded the statistical threshold.

**Figure 4.**
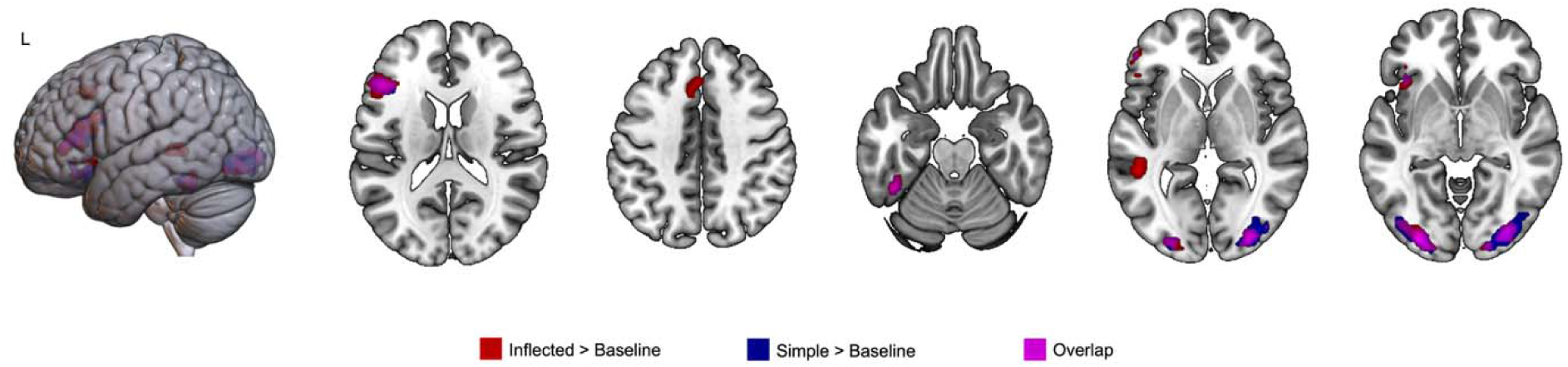
Whole-brain task activation for lexical processing. Brain activation maps depict voxels significantly engaged during word recognition in the whole-brain level (voxel-level threshold uncorrected *p* < .001, cluster extent threshold *k* _≥_ 30). Red = Inflected > Baseline, Blue = Simple > baseline, and Magenta = overlap of the two contrasts.

**Table 2.** MNI coordinates of significant activation identified in the whole-brain analysis.

| Brain Regions |  | Peak MNI coordinates | Z | Cluster Size |
| --- | --- | --- | --- | --- |
| *Word > Rest* |  |  |  |  |
| Lateral Occipital Cortex (inferior), Occipital Pole, Occipital Fusiform Gyrus | LH | [-30, -91, -4] | 8.2 | 2321 |
| Lateral Occipital Cortex (inferior), Occipital Pole | RH | [33, -88, -1] | 7.4 | 2002 |
| Frontal Orbital Cortex | LH | [-42, 26, -16] | 6.8 | 145 |
| Inferior Frontal Gyrus (pars triangularis, opercularis) | LH | [-48, 26, 17] | 6.5 | 1395 |
| Temporal Occipital Fusiform Cortex | LH | [-43, -49, -22] | 5.8 | 293 |
| *Inflected > Rest* |  |  |  |  |
| Lateral Occipital Cortex (inferior), Occipital Pole, Occipital Fusiform Gyrus | LH | [-30, -91, -4] | 7.6 | 2293 |
| Temporal Occipital Fusiform Cortex | LH | [-42, -46, -22] | 6.5 | 391 |
| Lateral Occipital Cortex (inferior) | RH | [33, -88, -1] | 6.4 | 1421 |
| Inferior Frontal Gyrus (pars triangularis, opercularis) | LH | [-48, 26, 17] | 6.3 | 1759 |
| Middle Temporal Gyrus (post) | LH | [-49, -43, 2] | 5.9 | 430 |
| Frontal Orbital Cortex | LH | [-40, 20, -7] | 5.4 | 240 |
| Paracingulate Gyrus | LH | [-3, 27, 42] | 5.2 | 347 |
| *Simple > Rest* |  |  |  |  |
| Lateral Occipital Cortex (inferior), Occipital Pole, Occipital Fusiform Gyrus | LH | [-30, -91, -4] | 7.7 | 2388 |
| Lateral Occipital Cortex (inferior), Occipital Pole | RH | [33, -88, -1] | 7.3 | 2732 |
| Frontal Orbital Cortex | LH | [-42, 26, -16] | 6.6 | 228 |
| Inferior Frontal Gyrus (pars triangularis, opercularis) | LH | [-48, 26, 15] | 6.3 | 1488 |
| Temporal Occipital Fusiform Cortex | LH | [-43, -49, -22] | 5.2 | 234 |

*Note.* Anatomical labels are from the Harvard–Oxford Cortical Atlas. All clusters survive whole-brain correction at voxel-level threshold of uncorrected $p < .001$ and cluster extent threshold of $k \geq 30$ . LH = left hemisphere; RH = right hemisphere.

When collapsed across all word conditions (*Word > Baseline*), robust activation was observed in a broad left-lateralized network associated with visual and lexical-semantic processing. Significant clusters were found in the left inferior frontal gyrus (pars triangularis and pars opercularis), frontal orbital cortex, temporal occipital fusiform cortex, and lateral occipital cortex (inferior division) bilaterally, with peak activity extending into the occipital pole.

The contrast Inflected > Rest revealed a largely overlapping activation pattern, including the left inferior frontal gyrus, middle and posterior temporal regions, temporal occipital fusiform cortex, frontal orbital cortex, and paracingulate gyrus. In addition, bilateral lateral occipital cortex (inferior division) showed significant engagement, reflecting visual processing demands.

The Simple > Rest contrast similarly activated the inferior frontal gyrus, frontal orbital cortex, and temporal occipital fusiform cortex, along with strong bilateral activation in the lateral occipital cortex and occipital pole. However, the extent of frontal and temporal involvement was more restricted compared to the inflected condition.

Notably, the direct contrast Inflected > Simple did not yield any suprathreshold clusters, suggesting that the two morphological conditions engaged similar neural systems when contrasted directly, despite observable differences in activation extent relative to baseline.

Together, these results indicate that both simple and inflected nouns recruited a left-lateralized fronto-temporo-occipital network, with no statistically significant differences in overall activation magnitude between conditions at the whole-brain level. However, when examined relative to baseline, inflected nouns additionally recruited middle and posterior temporal regions. This might reflect additional lexical-semantic processing demands associated with morphologically complex forms, which carry richer combinatorial meaning (e.g., case or topic information) than monomorphemic nouns.

### RSA results

The results of representational similarity analysis (RSA) are presented in **Fig. 5**. We assessed whether local multivoxel patterns reflected lexical frequency structure by correlating neural RDMs with theoretical RDMs based on either whole-word frequency (Zipf) or stem frequency (Zipf-Stem). RSA was conducted within six anatomically defined ROIs (IFGoper, IFGtri, AG, SMG, MTG, FuG). Group-level RSA effects were tested using one-sample t-tests (against zero) for each model (i.e., Zipf and Zipf-Stem), condition (All, Inflected, Simple), and ROI. To directly compare encoding strength across conditions, we additionally conducted paired two-sample t-tests between Inflected and Simple nouns. All *p*-values were corrected using false discovery rate (FDR) correction.

**Figure 5.**
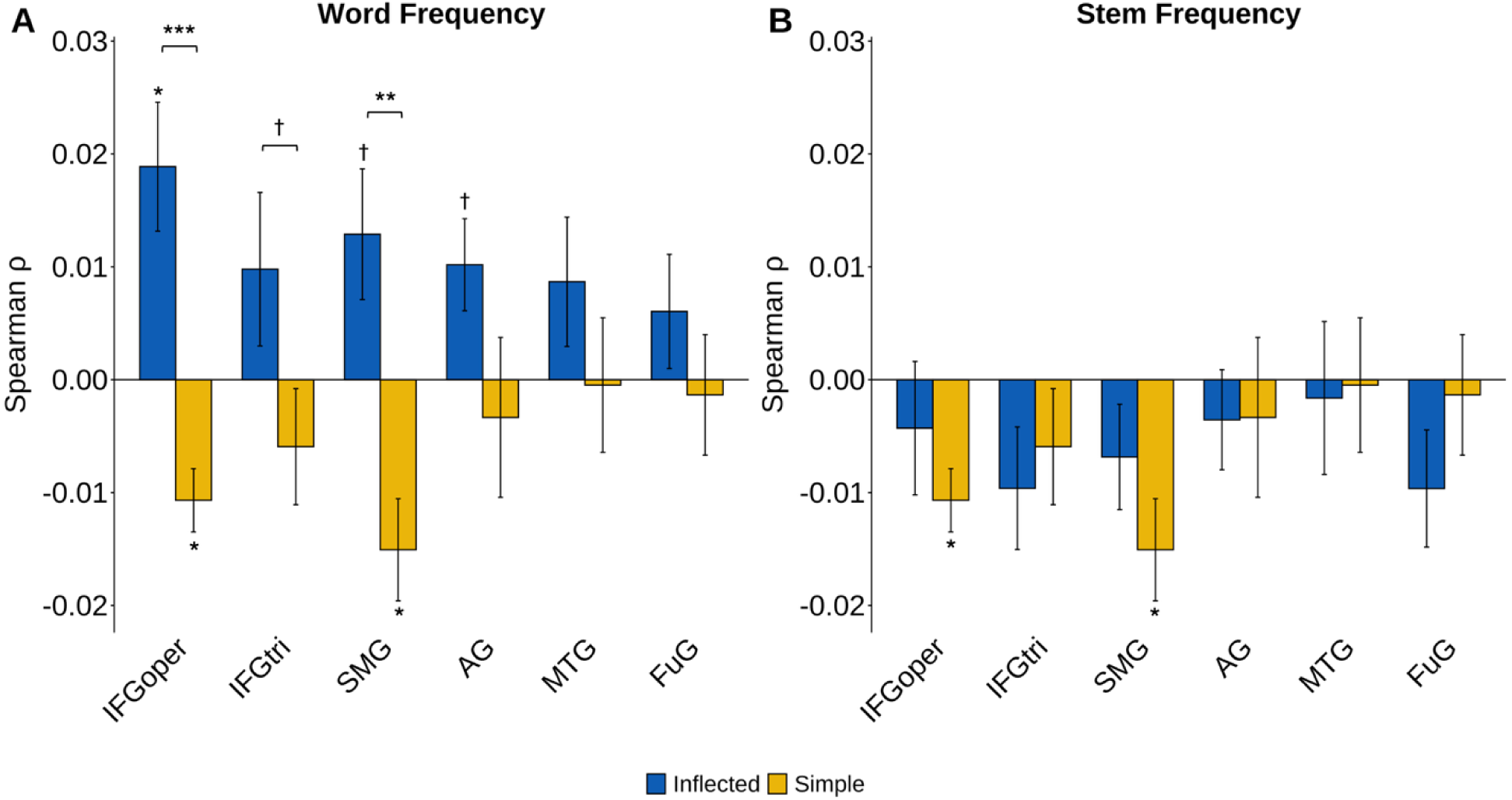
Representational similarity analysis (RSA) results. Mean Spearman *ρ* values between neural RDMs and (A) word-frequency (Zipf) or (B) stem-frequency (Zipf-Stem) model RDMs are plotted for each ROI (blue = Inflected; yellow = Simple); vertical bars represent ± SEM. Significance of each individual bar was tested with a one-sample t-test against zero and is indicated directly above the bar (* *p* < .05; ** *p* < .01; *** *p* < .001; † uncorrected *p* < .05, FDR-corrected within panel). Horizontal brackets linking the blue-yellow bar pairs denote paired-sample t-tests that compared Inflected and Simple correlations within the same ROI, using the same symbol conventions for significance.

In RSA using the word frequency (Zipf-word) model (**Fig. 5A**), several ROIs showed significant encoding for word frequency. One-sample *t*-tests revealed significant encoding of Zipf frequency for inflected nouns in IFGoper (*t*(20) = 3.30, *pFDR* = .025). Notably, AG (*t*(20) = 2.50, uncorrected *p* = .021) and SMG (*t*(20) = 2.22, uncorrected *p* = .038) also showed trend-level effects for inflected nouns, although these did not survive FDR correction. In contrast, RSA correlations for simple nouns significantly differed from zero in IFGoper (*t*(20) = −3.80, *pFDR* = .020) and SMG (*t*(20) = −3.33, *pFDR* = .025), indicating negative similarity effects, such that items with more similar surface frequency patterns produced more dissimilar neural patterns. Paired t-tests revealed significantly stronger encoding of word frequency for inflected compared to simple nouns in IFGoper (*t*(20) = 6.44, *pFDR* = .0001) and SMG (*t*(20) = 3.95, *pFDR* = .0096), suggesting that morphological complexity enhances the neural encoding of frequency information in these regions.

In contrast, RSA using the stem frequency model (Zipf-Stem; **Fig. 5B**) showed generally weaker effects. IFGoper (*t*(20) = −3.80, *pFDR* = .020) and SMG (*t*(20) = −3.33, *pFDR* = .025) showed significant negative RSA correlations for simple nouns. Trend-level negative RSA effects were observed in SMG for the All condition (*t*(20) = −2.41, uncorrected *p* = .026). However, no ROI showed significant above-zero encoding of Zipf-Stem values in the All or Inflected conditions, nor were significant differences found between inflected and simple nouns.

In sum, RSA revealed that whole-word frequency (Zipf) was significantly encoded in frontoparietal regions, especially IFGoper and SMG, with stronger encoding for inflected compared to simple nouns. Stem frequency (Zipf-Stem), in contrast, was only negatively encoded in IFGoper and SMG for simple nouns, suggesting that morphological complexity enhances sensitivity primarily to whole-word rather than stem-level frequency structure.

### Univariate analysis results

The results of the ROI analysis and illustration are presented in **Fig. 6**. We examined whether morphologically simple versus inflected nouns elicited differential activation across six predefined ROIs and whether this activation was modulated by surface-level (Zipf-word) (**Fig. 6B**) or stem-level (Zipf-stem) frequency (**Fig. 6C**).

**Figure 6.**
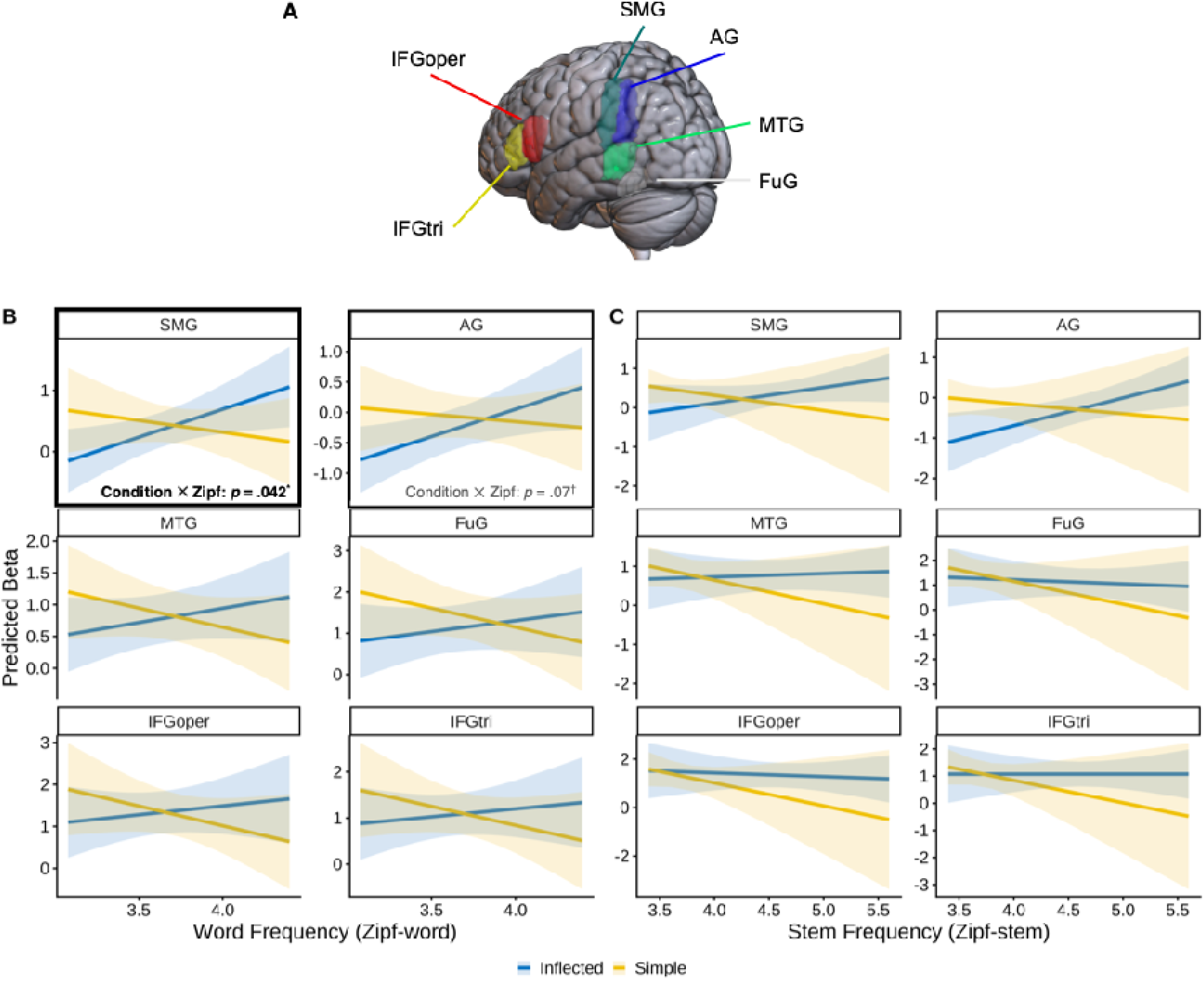
Regions of interest univariate analysis results. (A) Locations of anatomically defined ROIs: IFG pars opercularis (IFGoper), IFG pars triangularis (IFGtri), supramarginal gyrus (SMG), angular gyrus (AG), middle temporal gyrus (MTG), and fusiform gyrus (FuG). (B) Predicted BOLD β as a function of Zipf word frequency for inflected (blue) and simple (yellow) nouns, estimated from linear mixed-effects models. The Condition × Frequency interaction was significant in SMG (*p* = .042) and marginal in AG (*p* = .07). (C) The same analysis for Zipf stem frequency. Shaded ribbons represent 95% confidence intervals. \**p* < .05; ^†^*p* < .10.

Our primary analysis focused on Zipf frequency. We observed significant main effects of word frequency in both regions in the inferior parietal lobule (IPL), including the angular gyrus (*β* = 0.897, *SE* = 0.39, *t* = 2.3, *p* = .023) and supramarginal gyrus (*β* = 0.915, *SE* = 0.397, *t* = 2.3, *p* = .023), indicating increased activation for higher-frequency words. No other regions showed significant frequency effects after FDR correction.

Importantly, a significant interaction of Condition and Zipf-word emerged in the SMG (*β* = –1.30, *SE* = 0.634, *t* = –2.06, *p* = .042), reflecting stronger frequency effects for inflected compared to simple nouns. The AG also showed a marginal interaction (*β* = –1.14, *SE* = 0.624, *t* = –1.83, *p* = .07), although this did not survive correction. No significant main effects or interactions were found in IFG (pars opercularis/triangularis), or FuG.

We then tested the influence of stem frequency (Zipf-stem). A significant main effect was observed in the angular gyrus (*β* = 0.685, *SE* = 0.27, *t* = 2.54, *p* = .013), indicating greater activation for higher-frequency stems across conditions, though this effect did not remain significant after correction. No other regions showed reliable stem frequency effects or interactions with morphological condition. Importantly, SMG, which showed a robust interaction with surface frequency, did not exhibit significant modulation by stem frequency (*β* = –0.788, *SE* = 0.573, *t* = –1.37, *p* = .172).

Notably, as visible in **Figs. 6B–6C**, the highest frequency bin for simple nouns shows wider confidence intervals. This pattern reflects the lexical distribution of our stimuli set. As simple nouns equate surface and base frequency, they occupy a compressed frequency range and only a small number of items fall into the highest frequency bin (see **Fig. 1**). The reduced item variability in this range produces greater between-item variance and correspondingly wider intervals, likely reflecting the combination of a small number of high-frequency items and the inherently skewed frequency distributions observed in large lexical decision megastudies (Brysbaert et al., 2018; Keuleers et al., 2011; Yap & Balota, 2009). Importantly, this variance does not alter the overall pattern that surface-frequency effects are selectively and more robustly expressed for inflected nouns.

Together, these findings suggest that surface-level frequency exerts a stronger and more morphology-dependent influence on activation in parietal regions than stem-level frequency, consistent with a dissociation between surface-form and base-form frequency effects in morphological processing, in line with post-lexical accounts.

## DISCUSSION

The present study investigated how morphological complexity and lexical frequency jointly shape neural responses during visual word recognition, focusing on the dissociation between surface (whole-word) frequency and base (stem) frequency in Korean nominal inflection. Korean provides an interesting test case because its transparent agglutinative structure allows stem-based and surface-based statistics to be cleanly separated. By combining univariate mixed-effects modeling with representational similarity analysis (RSA) across six left-hemisphere language regions, we tested competing predictions of pre-lexical decomposition and post-lexical, whole-form access accounts. Three main findings were demonstrated. First, RSA revealed robust representational encoding of surface frequency in IFG pars opercularis (IFGoper) and supramarginal gyrus (SMG), with encoding significantly stronger for inflected than simple nouns. Second, base frequency did not show positive encoding in any ROI; instead, simple nouns produced negative RSA correlations with the base-frequency RDM in IFGoper and SMG. Third, univariate analyses in the inferior parietal lobule (IPL) revealed significant interactions of morphology and surface frequency, with activation increasing with surface frequency for inflected but not simple nouns. Collectively, these findings indicate that sustained neural activation patterns primarily reflect whole-form lexical statistics rather than stem-based morphological structure.

These results are in line with prior work in both Indo-European and Korean languages, while extending this literature by comparing surface- and stem-based frequency within a single inflectional system. Consistent with masked-priming and ERP evidence showing that high-frequency complex forms can be accessed more holistically (Kim et al., 2022; Lehtonen et al., 2007), our RSA and univariate findings demonstrate that surface frequency reliably modulates activation patterns in frontoparietal regions. Behavioral processing advantages for high-frequency inflected forms have been widely documented in lexical decision and naming tasks (Bertram et al., 2000; Kuperman et al., 2009), and our data converge with these findings by showing that surface frequency, but not stem frequency, shapes neural similarity and activation magnitude.

Neuroimaging studies have similarly implicated graded lexical-semantic and statistical influences on morphological processing. In languages such as English, Hebrew, Finnish, and Italian, fMRI work has shown that putative morphological effects often reduce to or interact with frequency, predictability, or semantic transparency (Bozic et al., 2013; Desai et al., 2006; Devlin et al., 2004). Our results extend this observation to Korean nominal inflection; despite its transparent morphological structure, the neural similarity in IFGoper and SMG was more strongly associated with whole-word frequency than with stem-based factors.

These findings refine our understanding of Korean morphological processing and situate it within broader cross-linguistic patterns. Prior fMRI research on Korean verb inflection has consistently shown overlapping activation for regular and irregular verbs in left IFG and temporal regions (Kim et al., 2024; Yim et al., 2006), suggesting that the processing of Korean inflectional morphology may not solely depend on categorical dichotomy. At the same time, these studies observed graded modulations by semantic homonymy and morphological complexity in occipitotemporal and angular gyrus regions (Kim et al., 2024), indicating sensitivity to experience-based properties. Our findings parallel this pattern in showing that although morphological complexity (simple vs. inflected) modulates representational strength, the representational structure was shaped primarily by surface-level lexical statistics.

The behavioral results complement the neural findings, further specifying how lexical statistics shape recognition of inflected forms. Both surface and base frequency facilitated lexical decisions, and neither interacted significantly with morphological complexity. This aligns with large lexical-decision megastudies showing that whole-word frequency is the primary predictor of behavioral performance (Keuleers et al., 2011; Yap & Balota, 2009; Yi et al., 2017). Importantly, the absence of a morphology × frequency interaction behaviorally parallels the pattern in the fusiform and MTG ROIs, which showed no reliable modulation by frequency.

The neural data, however, reveal distinctions that remain behaviorally silent. Whereas both frequency measures facilitate decision speed, only surface frequency shapes representational similarity in IFGoper and SMG, and only surface frequency interacts with morphology in parietal univariate activation. This dissociation suggests that behavioral measures and neural representational measures capture distinct aspects of word recognition; behavioral responses index the efficiency of the recognition and decision processes at the later stage of lexical access, whereas RSA and ROI activation reveal the internal representational architecture supporting this process. Inflected nouns with higher surface frequency may be accessed more efficiently behaviorally as they possess more stable or more distinct neural representations in IFGoper and IPL. Thus, the convergence between behavioral facilitation and neural representational structure strengthens the interpretation that surface frequency indexes a privileged pathway for accessing inflected forms.

A central theoretical question concerns what surface frequency encoding indeed represents neurally. In pre-lexical decomposition accounts, lexical access is assumed to proceed via early and obligatory segmentation into morphemic units, such that base frequency — reflecting the resting activation of stored stem representations — serves as a primary determinant of processing efficiency (Taft & Forster, 1975). Surface frequency, in this framework, is not expected to exert a dominant influence during initial processing stages; rather, its effects would emerge only at later stages where decomposed morphemes are recombined and matched against whole-word representations. In contrast, post-lexical or full-form access accounts hold that surface frequency directly indexes the strength of stored whole-word lexical entries, with higher-frequency forms benefiting from more robust representations and thus more efficient retrieval. Two aspects of our findings support the latter interpretation.

First, when examined within the inflected-noun stimuli, surface and base frequency demonstrated only moderate correlation and produced clearly distinct representational geometries (see **Fig. S1**). If surface-frequency effects were driven by stem-level statistics, one would expect base frequency to exhibit parallel RSA encoding, which it did not. Second, surface-frequency encoding localized primarily to IFGoper and SMG, regions implicated in lexical retrieval, semantic integration, and competition resolution (Binder et al., 2009; Lau et al., 2008; Thompson-Schill et al., 1997), rather than in occipitotemporal regions, which has been functionally linked to early morpho-orthographic segmentation (Gold & Rastle, 2007; Solomyak & Marantz, 2010). This spatial dissociation implies that the RSA patterns reflect downstream lexical-semantic processes tied to whole-form access rather than sublexical parsing.

Another interpretation can be derived from the Augmented Addressed Morphology model (AAM; Caramazza et al., 1988) proposes that lexical access can proceed via both whole-word representations and morphological decomposition, with the relative contribution of each route depending on factors such as frequency and representation strength. Under this framework, high-frequency inflected forms are preferentially accessed via whole-word representations, whereas lower-frequency forms are more likely to engage decomposition processes. From this perspective, the surface-frequency effects observed in IFGoper and SMG may reflect the degree to which specific inflected forms have developed robust whole-word representations through repeated exposure. Inflected nouns with high surface frequency may function as independent lexical entries whose representational geometry is shaped by cumulative experience, consistent with the AAM’s prediction that frequency modulates the relative efficiency of the whole-word access route. This interpretation aligns with experience-based and connectionist accounts in which lexical representations emerge from statistical regularities in form–meaning mappings rather than from strictly compositional morpheme-based storage (Baayen et al., 2011; Bybee, 1995; Plaut & Gonnerman, 2000).

Region-specific patterns further refine understanding of the neural architecture supporting morphological processing. IFG pars opercularis showed robust RSA encoding of surface frequency for inflected nouns, suggesting that this region participates in representing graded lexical statistics rather than merely implementing competitive selection or morpho-syntactic parsing as traditionally assumed. This finding aligns with work linking posterior IFG to the integration of form and meaning under conditions of lexical competition or ambiguity (Thompson-Schill et al., 1997; Whitney, Jefferies, et al., 2011).

In the IPL, both SMG and AG showed univariate interactions of morphology and surface frequency, with increasing activation for higher-frequency inflected forms. IPL has long been associated with lexical–semantic retrieval and the integration of distributed semantic features (Binder et al., 2009; Price et al., 2015), as well as with controlled semantic access in tasks requiring selection or feature weighting (Graves et al., 2010; Whitney, Jefferies, et al., 2011). Our findings reinforce this functional characterization by indicating that IPL encodes the accessibility or retrieval strength of whole forms rather than engaging in decomposition of stems.

In contrast, MTG showed weak or inconsistent frequency effects. Given its role in semantic control and relational processing (Davey et al., 2016; Whitney, Kirk, et al., 2011), MTG activation may emerge more strongly in tasks with greater semantic integration demands such as semantic judgment rather than in lexical decision. Additionally, although whole-brain analyses identified fusiform activation for all word stimuli, our anatomically defined fusiform ROI did not show reliable frequency modulation. This likely reflects the broad mask used, which spans subregions with distinct functional profiles. Only mid-fusiform cortex reliably shows graded frequency and lexicality effects; more precise functional ROIs might reveal effects not visible here.

The present findings offer support for whole-word access accounts over obligatory decomposition models, which predict robust stem-frequency effects and decomposition signatures in all morphologically complex words (Rastle et al., 2004; Taft & Forster, 1975). Although early ERP studies support rapid form-based segmentation, such processes may be transient and not reflected in sustained BOLD patterns. Our results instead align with flexible or hybrid models in which whole-word access pathways are strengthened by lexical experience and decomposition is engaged selectively when whole-form access is inefficient (Baayen et al., 2011; Milin et al., 2017).

The results are also compatible with the broader view that morphological computation is distributed and usage-dependent rather than categorical and rule-governed. For Korean nominal inflection, surface frequency shapes both representational structure and activation dynamics, suggesting that frequent inflected forms may become entrenched as distinct lexical representations. This interpretation aligns with usage-based and connectionist perspectives in which morphological structure emerges from learned statistical regularities (Bybee & Hopper, 2001; Plaut & Gonnerman, 2000), as well as with the AAM framework, which predicts that high-frequency inflected forms can be accessed via a direct whole-word route, effectively functioning as independent lexical entries (Caramazza et al., 1988).

Some limitations should be acknowledged. First, temporal resolution of fMRI limits direct observation of rapid pre-lexical segmentation processes occurring within the first approximately 200 ms of word processing, the processing stage indexed by masked priming and early ERP components. Early form-based segmentation may nonetheless occur transiently and be superseded by the whole-form access patterns visible in sustained BOLD responses. Multimodal recordings combining EEG/MEG with fMRI would be necessary to adjudicate whether pre- and post-lexical decomposition pathways operate sequentially during inflected word recognition.

Relatedly, stem syllable counts necessarily differ between simple and inflected conditions due to the addition of a monosyllabic suffix in inflected forms. Although total word length was matched across conditions and first-syllable frequency did not differ, we cannot entirely rule out that differences in stem length contributed to the observed morphology × frequency interactions. However, if stem length were driving the effects, one would expect base (stem) frequency—which directly indexes stem-level processing—to show stronger neural encoding for inflected nouns with shorter stems, which was not observed.

Additionally, our sample size (*N* = 21) and anatomically defined ROI approach afford adequate power for medium-to-large effects but may miss subtle regional interactions, particularly in heterogeneous regions such as MTG and mid-fusiform cortex. Similarly, the lexical-decision task emphasizes decisional speed and accuracy rather than automatic comprehension; converging evidence from speeded naming, semantic judgment, or passive reading tasks would strengthen claims about the generality of surface-frequency effects.

## CONCLUSION

In conclusion, the present findings provide the first fMRI evidence comparing surface and base frequency effects on neural representational structure in morphologically complex Korean words. The results demonstrate that morphological complexity modulates neural encoding of surface rather than base frequency and that morphological condition does not yield discrete category representations in classic language areas. These findings argue against purely decompositional theories and instead support distributional or hybrid models in which both decomposition and whole-form access routes exist, with their engagement determined by lexical statistics such as surface frequency. Rather than reflecting an obligatory parsing process, morphological processing emerges from dynamic interplay between a word’s structure and its statistical properties, shaped by both form-based and meaning-based constraints (Weissbart & Martin, 2024).

