## Supplementary material for "Neural Sensitivity to Word Frequency Modulated by Morphological Structure: Univariate and Multivariate fMRI Evidence from Morphologically Complex Words": Fig. S1

### **Supplementary Materials**

**Stimulus Frequency Validation**

We validated that word and stem frequency measures exhibit distinct distributional properties despite moderate item-level correlation. Beyond item-level properties, we assessed whether the two frequency measures organize the stimulus space differently for inflected nouns using representational similarity analysis (RSA). Representational dissimilarity matrices (RDMs) were constructed by computing absolute pairwise frequency differences for all inflected word pairs (see Methods). Although word and stem frequency showed moderate item-level correlation for inflected words (Pearson *r* = .60, *p* < .001; **Fig. S1A**), their RDM structures were largely independent (Spearman *ρ* = .24, *p* < .001; **Fig. S1C**), accounting for only 5.7% shared variance (*ρ²* = .057). This 94.3% independence indicates that the two measures create fundamentally different representational geometries for inflected nouns, as visualized in the RDM difference map (**Fig. S1B**). Together, these analyses demonstrate that moderate item-level correlation does not preclude independent representational structures when distributional properties differ substantially. The distinct variance structures and representational geometries enabled subsequent neural analyses to dissociate word-level and stem-level frequency effects.

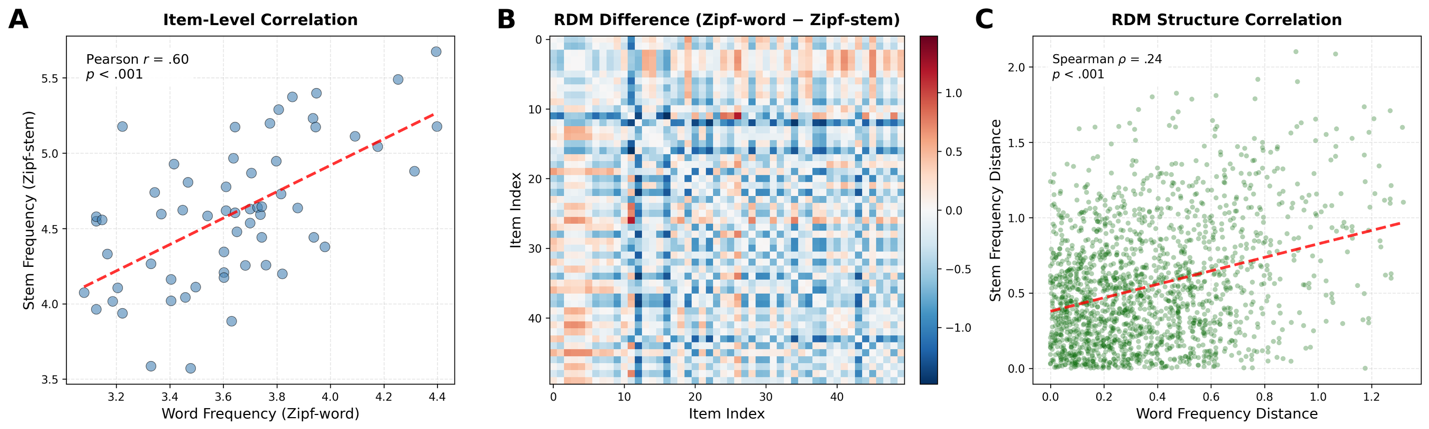

**Supplementary Figure S1.** Relationship between word and stem frequency measures for inflected nouns. (A) Item-level correlation between surface frequency and base frequency (N = 60). Each point represents one inflected noun; red dashed line indicates linear regression fit. (B) Representational dissimilarity matrix (RDM) difference map computed as Surface RDM minus Base RDM. Blue regions indicate greater dissimilarity in surface frequency; red regions indicate greater dissimilarity in base frequency. (C) Structure-level correlation between surface and base RDMs across all pairwise word comparisons (N = 3,540 pairs). Each point represents one word pair; red dashed line indicates monotonic relationship. Green points show the distribution of pairwise frequency distances.
